# Resistance patterns in drug-adapted cancer cell lines reflect the complex evolution in clinical tumours

**DOI:** 10.1101/2024.01.20.576412

**Authors:** Helen E. Grimsley, Magdalena Antczak, Ian G. Reddin, Katie-May McLaughlin, Andrea Nist, Marco Mernberger, Thorsten Stiewe, Tim R. Fenton, Daniel Speidel, Catherine Harper-Wynne, Karina Cox, Jindrich Cinatl, Mark N. Wass, Michelle D. Garrett, Martin Michaelis

## Abstract

**Background:** Here, we introduce a novel set of triple-negative breast cancer (TNBC) cell lines consisting of MDA-MB-468, HCC38, and HCC1806 and their sublines adapted to cisplatin, doxorubicin, eribulin, paclitaxel, gemcitabine, or 5-fluorouracil.

**Methods:** The cell lines were characterized by whole exome sequencing and the determination of drug-response profiles. Moreover, genes harbouring resistance-associated mutations were investigated using TCGA data for potential clinical relevance.

**Result:** Sequencing combined with TCGA-derived patient data resulted in the identification of 682 biomarker candidates in the pan-cancer analysis. Thirty-five genes were considered the most promising candidates because they harboured resistance-associated variants in at least two resistant sublines, and their expression correlated with TNBC patient survival. Exome sequencing and response profiles to cytotoxic drugs and DNA damage response inhibitors identified revealed remarkably little overlap between the resistant sublines, suggesting that each resistance formation process follows a unique route. All of the drug-resistant TNBC sublines remained sensitive or even displayed collateral sensitivity to a range of tested compounds. Cross-resistance levels were lowest for the CHK2 inhibitor CCT241533, the PLK1 inhibitor SBE13, and the RAD51 recombinase inhibitor B02, suggesting that CHK2, PLK1, and RAD51 are potential drug targets for therapy-refractory TNBC.

**Conclusions:** We present novel preclinical models of acquired drug resistance in TNBC and many novel candidate biomarkers for further investigation. The finding that each cancer cell line adaptation process follows an unpredictable route reflects recent findings on cancer cell evolution in patients, supporting the relevance of drug-adapted cancer cell lines as preclinical models of acquired resistance.

## Introduction

Triple-negative breast cancer (TNBC) is characterized by the absence of estrogen, progesterone, and HER2 receptors ^1^. TNBC is responsible for approximately 15% of breast cancer cases and is associated with a poorer prognosis than hormone receptor- or HER2-positive breast cancers ^1,2^. Current TNBC therapies are largely based on cytotoxic anticancer drugs, including platinum drugs, anthracyclins, eribulin, gemcitabine, paclitaxel, and 5-fluorouracil ^1^. TNBC often responds well initially to cytotoxic chemotherapy, but recurrence and resistance are common, eventually leading to therapy failure. This combination of an initial high response rate followed by rapid resistance is referred to as the ‘TNBC paradox’ ^1,3^. To improve TNBC therapy outcomes, new treatment approaches are needed, particularly those that are effective against treatment-refractory disease characterized by acquired resistance to cytotoxic chemotherapy.

In contrast to intrinsic drug resistance (which occurs independently of therapy and is a consequence of pre-existing often stochastic events in cancer cells), acquired resistance is the direct consequence of selection and adaptation processes caused by cancer treatment (directed tumor evolution) ^4–8^. Understanding acquired resistance mechanisms is essential for optimizing cancer treatment for patients with therapy-refractory tumors.

Drug-adapted cancer cell lines are preclinical models that have been shown to reflect clinically relevant acquired drug resistance mechanisms in numerous studies ^4,9–17^. Furthermore, drug-adapted cell lines enable detailed functional and systems-level studies that are not possible using clinical samples ^4^.

Here, we introduce a novel set of three parental TNBC cell lines and their 15 sublines adapted to cisplatin, doxorubicin, eribulin, gemcitabine, paclitaxel, or 5-fluorouracil. These cell lines were characterized by whole exome sequencing and the determination of response profiles to cytotoxic anti-cancer drugs and a panel of DNA damage repair inhibitors. The resulting data showed that each resistance formation process follows an individual and unpredictable route. The combined analysis of resistance-associated mutations in combination with patient data from The Cancer Genome Atlas (TCGA) ^18^ identified 35 novel candidate resistance biomarkers for further investigation.

## Results

### Project cell line panel

Here, we characterized a cell line panel consisting of the parental TNBC cell lines MDA-MB-468, HCC38, and HCC1806 and their sublines adapted to grow in the presence of cisplatin, doxorubicin, eribulin, paclitaxel, gemcitabine, or 5-fluorouracil, which are all drugs used for the treatment of TNBC (Fig. 1A, Suppl. File 1) ^19–25^. The drug-resistant sublines were established by continuous exposure to stepwise increasing drug concentrations as previously described ^16^. All parental cell lines were initially sensitive to therapeutic concentrations of the respective drugs, as indicated by IC_50_ (concentration that reduces cell viability by 50%) values within the range of clinical drug plasma concentrations (C_max_) (Suppl. Fig. 1, Suppl. File 1) ^26^. The relative resistance factors (IC_50_ drug-adapted subline/ IC_50_ respective parental cell line) ranged from 5.5-fold (HCC38^r^PCL^2.5^) to 5916.7-fold (HCC1806^r^ERI^50^) (Fig. 1B, Suppl. File 1).

**Figure 1.**
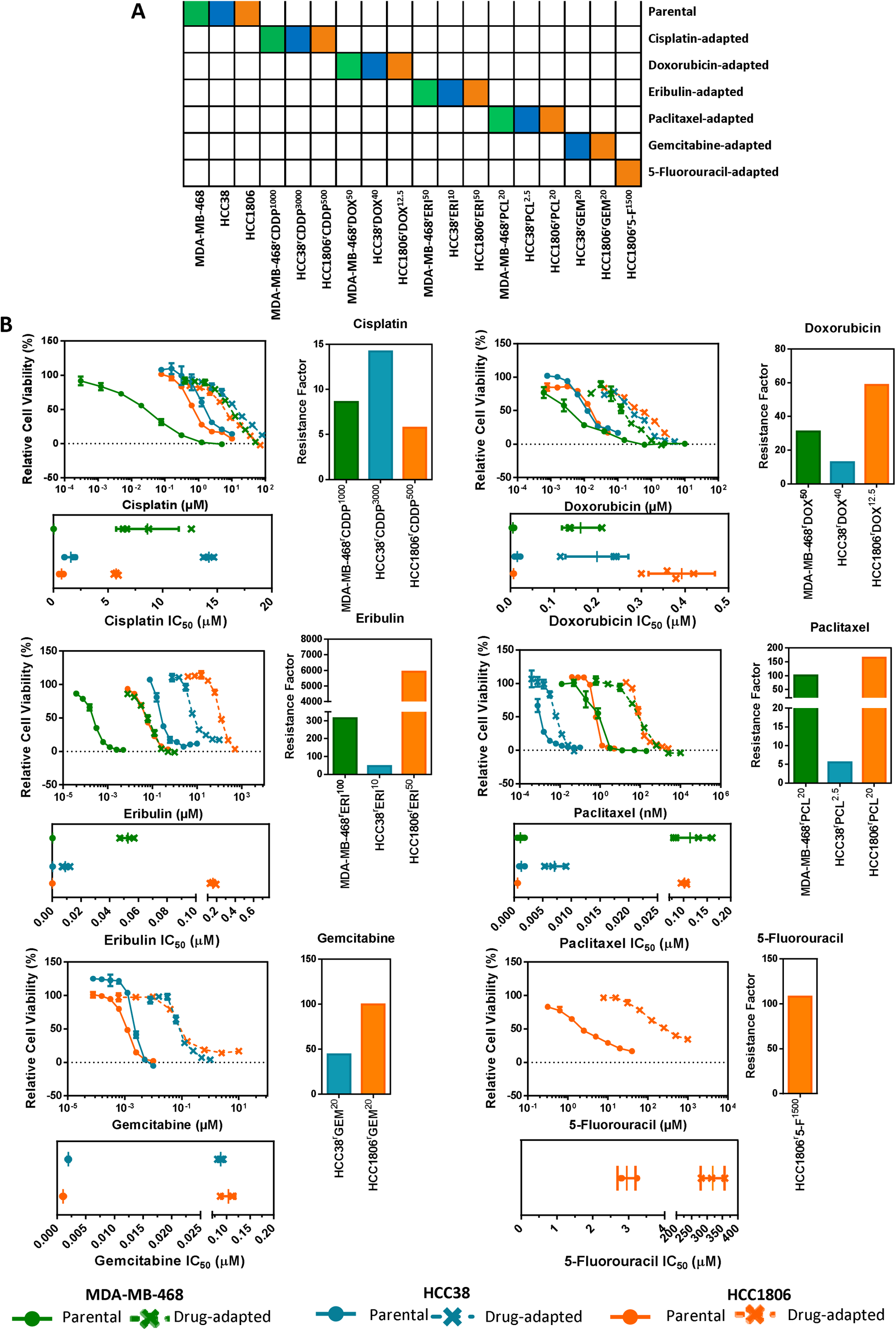
Confirmation of the resistance status of the project cell lines. A) Panel of drug-naïve (MDA-MB-468, HCC38, HCC1806) and drug-adapted Triple Negative Breast Cancer cell lines. B) Left: dose–response curve; bottom: IC_50_ values; right: resistance factor (IC_50,_ drug-adapted subline/IC_50,_ respective parental cell line)); when drug-naïve and drug-adapted cell lines are treated with the respective agent: cisplatin, doxorubicin, eribulin, paclitaxel, gemcitabine, or 5-fluorouracil. Circles indicate drug-naïve cell lines, and crosses indicate drug-adapted cell lines. Green, MDA-MB-468-derived; blue, HCC38-derived; orange, HCC1806-derived. The data are from ≥ 3 independent experiments, and the statistics were calculated using Student’s t-test and are plotted as the means ± SDs.

### Characterization of the cell line panel by whole exome sequencing

The cell line panel was investigated by whole exome sequencing. Among the identified variants, missense variants were most common, followed by synonymous variants (Suppl. Fig. 2A). Insertions/deletions (INDELs), frameshift mutations, stop-gain, stop-loss, and splice variants were identified at lower frequencies (Suppl. Fig. 2A). Between 217 (HCC38^r^DOX^40^) and 952 (HCC38^r^GEM^20^) variants differed in the drug-adapted sublines relative to the respective parental cell lines (Suppl. Fig. 2B).

We grouped the resistance-associated variants into five categories (Fig. 2A, see methods): 1. *Gained variants*, variants only called in the drug-adapted subline but detectable at low confidence in the respective parental cell line; 2. *De novo variants*, variants called in the drug-adapted subline but undetectable in the respective parental cell line; 3. *Not-called variants*, variants only called in the parental cell line but detectable with low confidence in the resistant subline; 4. *Lost variants*; variants called in the parental cell line but undetectable in the drug-adapted subline; and 5. *Shared variants*; variants called in both the parental and the respective drug-adapted sublines (Fig. 2A).

**Figure 2.**
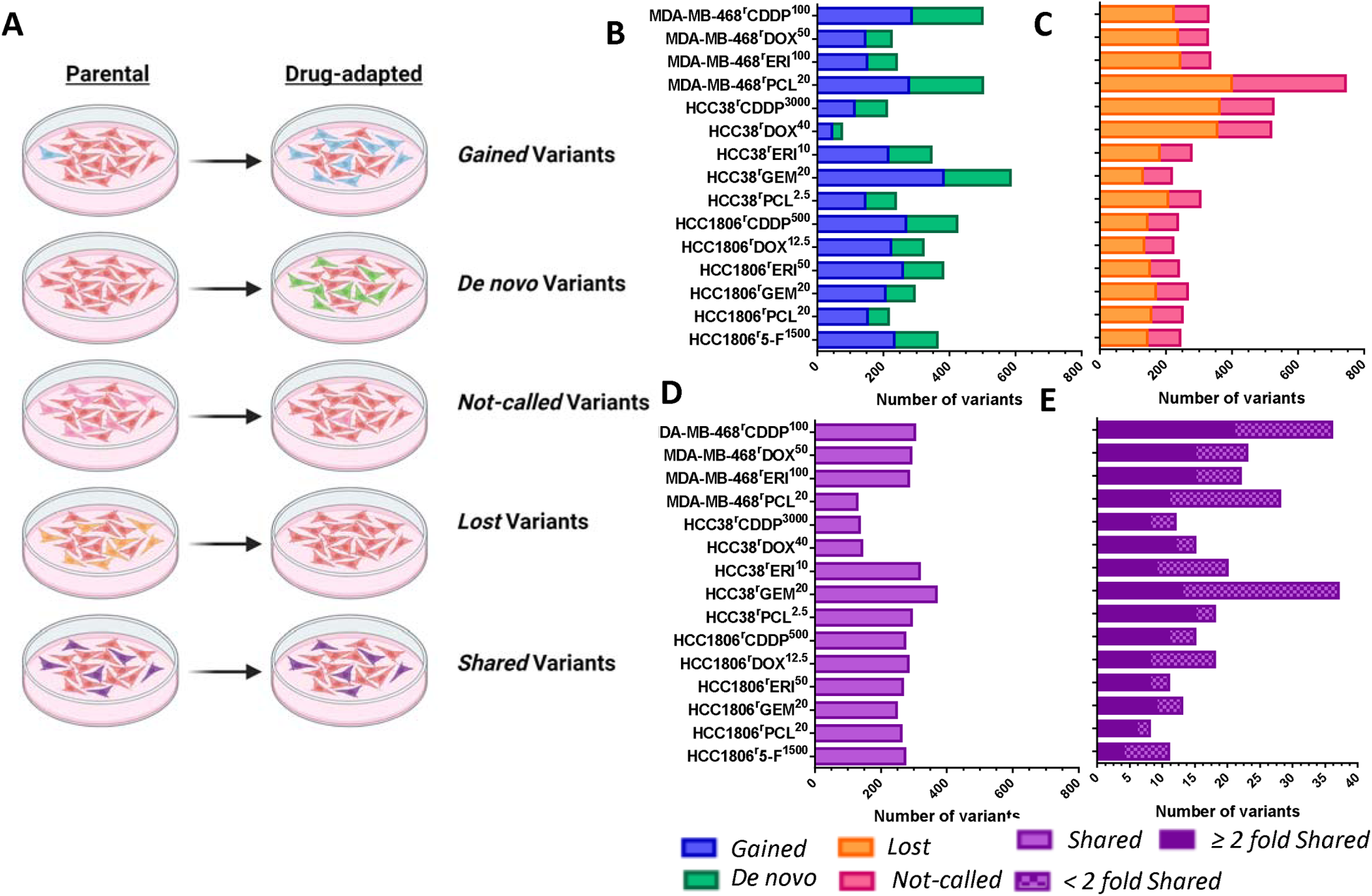
Genomic characterization of drug-adapted cell lines. A) Diagram illustrating the differences between *gained, de novo*, *not called*, *lost* and *shared* variants. B) Count of *Gained* (blue) and *De novo* (green) variants, C) count of *Lost* (orange) and *Not-called* (pink) variants, D) left panel; count of all *Shared* (purple) variants, right panel; two-fold increase or decrease of shared variants.

The number of *gained* variants ranged from 44 (HCC38^r^DOX^40^) to 381 (HCC38^r^GEM^20^), the number of *de novo* variants ranged from 31 (HCC38^r^DOX^40^) to 225 (MDA-MB-468^r^PCL^20^), the number of *not-called* variants ranged from 88 (HCC38^r^GEM^20^ and HCC1806^r^DOX^12.5^) to 345 (MDA-MB-468^r^PCL^20^), and the number of *lost* variants ranged from 129 (HCC38^r^GEM^20^) to 398 (MDA-MB-468^r^PCL^20^) (Fig. 2B, Fig. 2C, Suppl. File.2 and 3). The number of *shared* variants that were both called in the parental cell lines and their sublines ranged from 128 (MDA-MB-468^r^PCL^20^) to 368 (HCC38^r^GEM^20^) (Fig. 2D, Suppl. File 2 and 3). The number of *shared* variants that increased by at least two-fold in the resistant sub-lines vs. the respective parental ranged from four (HCC1806^r^5-F^1500^) to 21 (MDA-MB-468^r^CDDP^1000^), whilst the number of shared variants that decreased by at least two-fold ranged from two (MDA-MB-468^r^PCL^20^) to 24 (HCC38^r^GEM^20^) (Fig. 2E, Suppl. File 2).

### Analysis of the distribution of *de novo* variants

To identify variants that may have a functional role in drug resistance, we initially considered the 81 genes that harbored *de novo* variants in at least two different sublines from more than one parental cell line (Fig. 3A, Suppl. File 4). This list included 46 genes that have already been reported to be involved in drug resistance in cancer and 33 new candidate genes with a possible role in drug resistance (Fig. 3A, Suppl. File 4). Notably, 24 of the 33 new candidate genes are reported to be relevant in cancer (Fig. 3A, Suppl. File 4).

**Figure 3.**
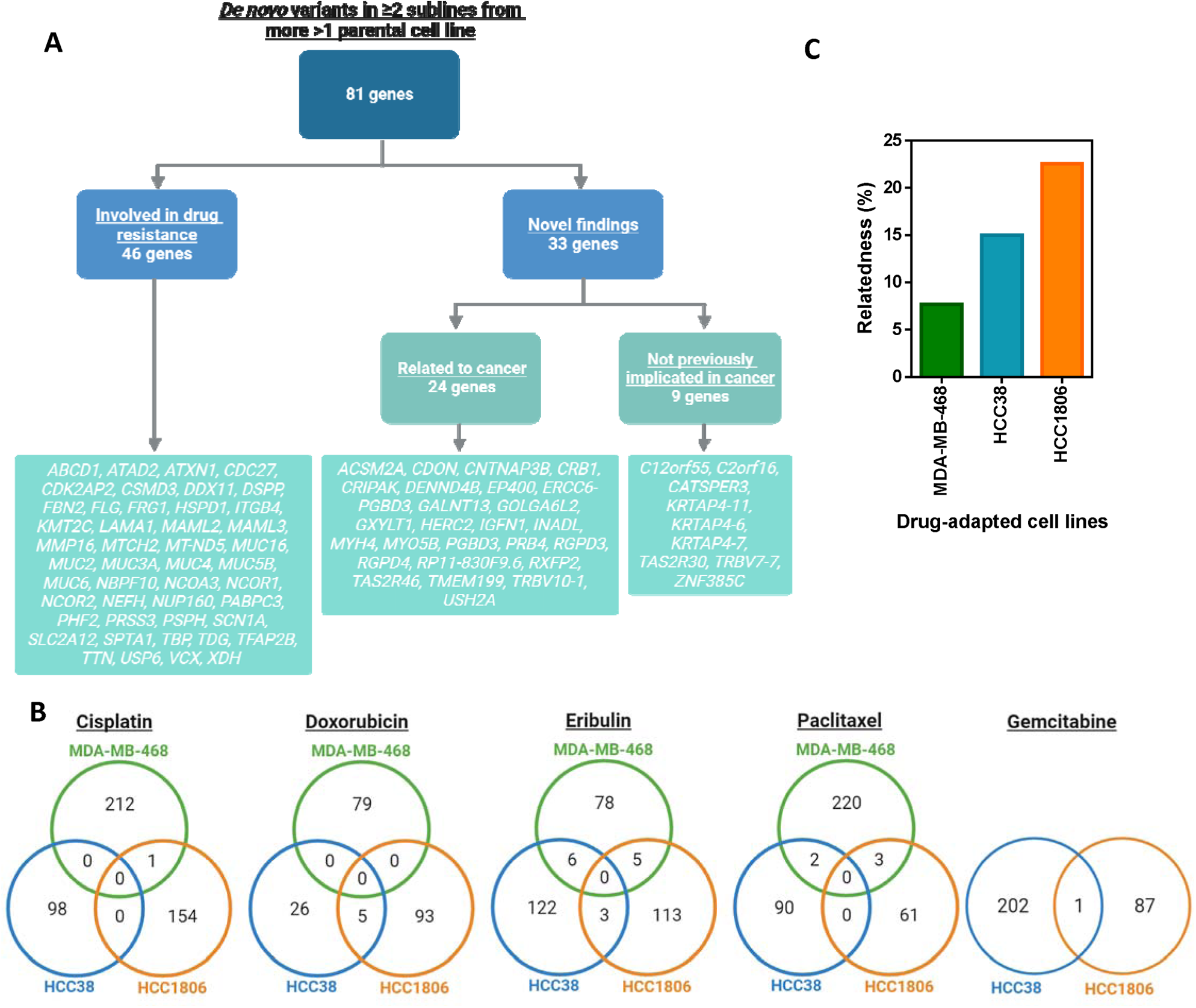
Identification of novel candidates associated with therapy failure. A) Flow chart of genes with *de novo* variants observed in two or more sublines from more than one parental cell line. B) Venn diagrams of *de novo* variants shared between sublines adapted to the same drug. C) Summary of relatedness between sublines drug-adapted from the same parental cell line (%).

Four of the five genes with the greatest number of *de novo* variants in the drug-adapted sublines were mucin (MUC) genes. *MUC6* had *de novo* variants in 15 sublines, *MUC2* in 14 sublines, *MUC4* in 13 sublines, and MUC16 in nine sublines (Fig. 3A, Suppl. File 4). The MUC genes are large genes that are known to be commonly mutated in cancer and have been reported to be involved in cancer cell drug resistance ^27–31^. *De novo* mutations in CDC27, which has also been linked to drug resistance in cancer, were also detected in nine drug-resistant sublines ^32,33^ (Fig. 3A, Suppl. File 4).

*GXYLT1*, *KRTAP4-11*, and *RGPD4* were amongst those genes, which had not previously been associated with drug resistance in cancer that displayed *de novo* mutations in a high number (7) of drug-resistant sublines (Fig. 3A, Suppl. File 4). A GXYLT1 mutation promoted metastasis in colorectal cancer through MAPK signalling, a pathway known to confer resistance to a range of anti-cancer drugs ^34–37^. *RGPD4* mutations are correlated with vascular invasion in HBV-associated hepatocellular carcinoma, and it is known that there is an overlap between pro-angiogenic, pro-metastatic, and resistance-associated signalling in cancer ^35,38^. There is no known link between KRTAP4-11 and cancer, but *KRTAP4-11* expression levels have been reported to predict the methotrexate response in rheumatoid arthritis patients ^38^. Hence, it seems plausible that the products of these genes may be involved in cancer cell drug resistance.

Taken together, our analysis identified 48 genes known to be involved in cancer cell drug resistance alongside 33 novel candidates potentially contributing to therapy failure. Further research will be required to characterize the roles of these individual genes in detail.

When we compared the overlaps between *de novo* variants shared between sublines adapted to the same drug, the numbers were too small to draw any meaningful conclusions (Fig. 3B, Suppl. Fig. 3A).

Notably, *de novo* variants in drug-resistant sublines may not always represent actual novel variants that are selected because they contribute to cancer cell resistance. Many apparent *de novo* mutations may have already been present in a small fraction of the cells of the parental cell line but may not have been detected due to the sequencing depth. Hence, overlaps in *de novo* variants between sublines of the same parental cell line can also be used to indicate the levels of relatedness between the founding subpopulations of the different resistant sublines.

Analysis of the *de novo* variants shared between the sublines from the same parental cell line indicated the largest overlap. On average, there was a 22.6% overlap among the HCC1806 sublines, followed by a 15.0% overlap among the HCC38 sublines and a 7.7% overlap among the MDA-MB-468 sublines (Fig. 3C). However, there were also noticeable differences in the overlaps between *de novo* variants identified in each of the sublines from the same parental cell line. For example, only three *de novo* variants were shared between HCC38^r^CDDP^3000^ (out of 98 in total, 3.1%) and HCC38^r^PCL^2.5^ (out of 92 in total, 3.3%), while 53 variants were shared between HCC38^r^ERI^10^ (out of 131 in total, 40.5%) and HCC38^r^GEM^20^ (out of 203 in total, 26.1%) (Fig. 3C, Suppl. Fig. 3B). These numbers suggest that there are no pre-existing cell line subpopulations that are consistently selected in response to anti-cancer drug treatment.

### Protein functions related to variants that changed in drug-resistant sublines

Next, we used the Gene Ontology (GO) annotation to perform an analysis of the protein functions associated with genes present in the *de novo*, *gained*, *not called*, and *lost* variant sets as well as *shared* variants with a two-fold increase or decrease in allele frequency (Suppl. Fig 4A, B).

There was limited overlap between the GO terms for the variants detected in the sublines adapted to the same drug (Suppl. Fig. 4C, E). The extracellular matrix-related GO terms ‘extracellular matrix constituent lubricant activity’, ‘extracellular matrix’, and ‘maintenance of gastrointestinal epithelium’ were most common, which reflects the high number of variants observed in the mucin genes (Suppl. Fig. 4C, E).

GO term analysis of the sublines from the same parental cell line revealed very similar results, again revealing an overrepresentation of extracellular matrix-related GO terms (Suppl. Fig. 4D, F). Further research will be required to investigate the potential role of mucins and the extracellular matrix in acquired drug resistance in TNBC cells.

### Potential clinical relevance of selected variants

The potential clinical relevance of genes harboring *de novo*, *gained*, and *shared* variants with a two-fold increase in allele frequency in the resistant subline as well as genes harboring truncating variants was analysed using patient data derived from The Cancer Genome Atlas (TCGA) ^39^. Notably, there were only data available from patients treated with cisplatin, doxorubicin, gemcitabine, paclitaxel, and/or 5-fluorouracil, but no data on eribulin treatment were available.

We performed two analyses, one pan-cancer analysis, in which we considered all patient survival data available for the drugs, and a second analysis, in which we considered TNBC patients and for which only doxorubicin and paclitaxel data were available (Fig. 4A). The pan-cancer analysis included data from 29 TCGA cancer types for which mutation status and gene expression data were available (Fig 4B).

**Figure 4.**
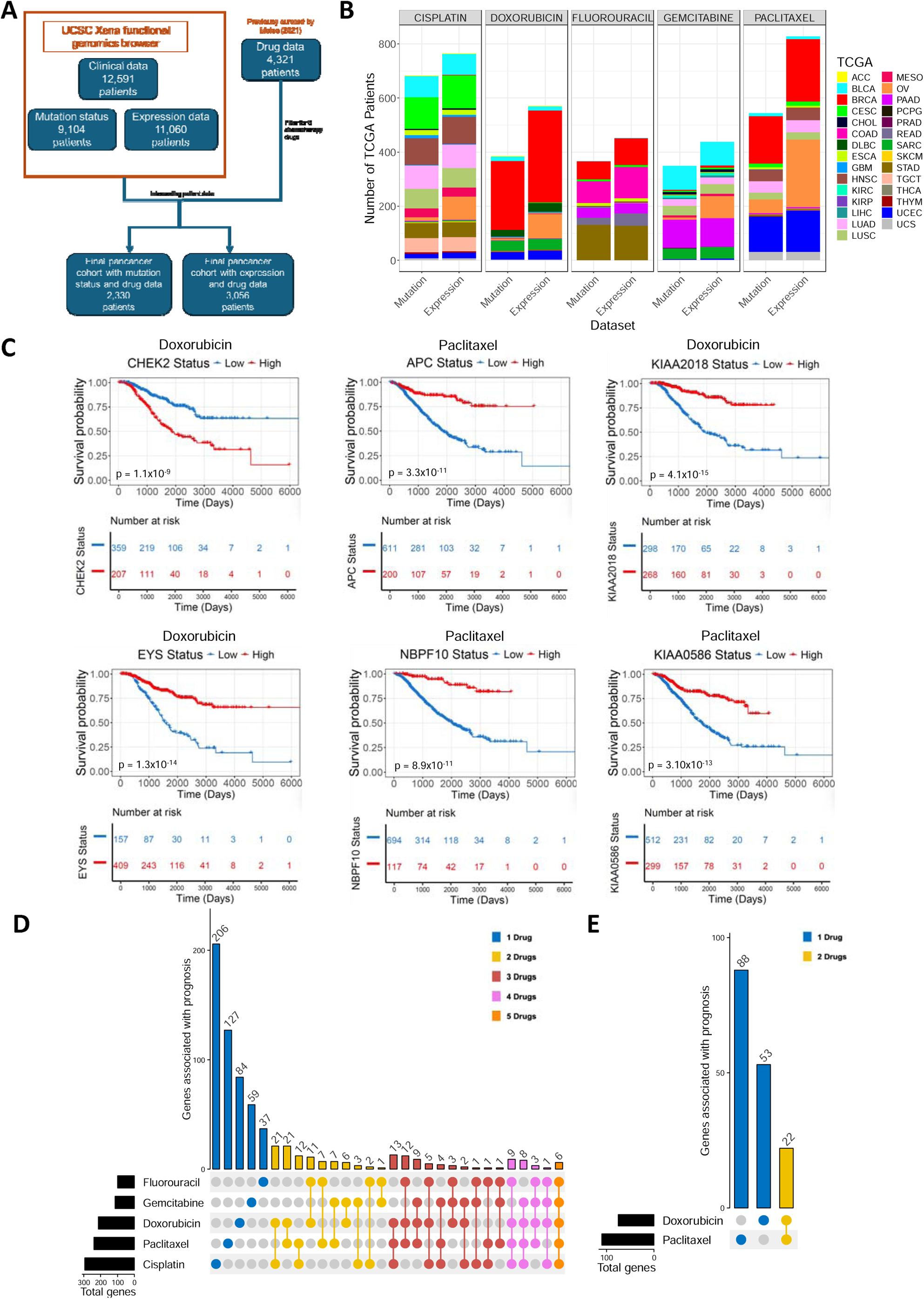
Tumor patient data available for mutations in resistance-associated genes. (A) TCGA pan-cancer datasets for mutation status and gene expression. Only patients for which clinical, drug and mutation status/gene expression data was available for were considered in the TCGA pan-cancer analysis. (B) TCGA pan-cancer mutation status and expression data available for chemotherapy drugs for 29 TCGA cancer classifications. (C) Kaplan-Meier plots for gene expression with most significant association with prognosis in the pan-cancer dataset. Log-rank test was the statistical test used with multiple test correction performed using Benjamini-Hochberg method. (D-E) Genes for which expression is significantly associated with patient prognosis. Upset plot showing the number of genes that are associated with patient prognosis for (D) pan-cancer and (E) TNBC.

Six cases with at least two mutations in a resistance-associated gene were associated with patient prognosis (Suppl. File 5). As the number of resistance-associated genes with mutations in patients was low, we also considered gene expression status associations with prognosis. For 1,018 cases there was a significant association between gene expression and patient prognosis. This included genes, whose products were known to play a role in cancer cell drug resistance, such as CHEK2 ^40–42^ and APC ^43–45^ (Fig. 4C, Suppl. File 5). Moreover, we also identified novel candidates, which had not previously been suggested to be involved in cancer cell drug resistance, including KIAA2018, EYS, NBPF10, and KIAA0586 (Fig. 4C, Suppl. File 5).

We further determined the association of the expression of genes harboring *de novo*, *gained*, and *shared* variants with a two-fold increase as well as genes harboring truncating variants with patient survival in response to treatment with cisplatin, doxorubicin, gemcitabine, paclitaxel, and 5-fluorouracil (Fig. 4 D-E, Suppl. File 5). In total, the expression of 682 genes was significantly correlated with patient survival in response to at least one drug in the pan-cancer data. For 513 of these 682 genes, gene expression was associated with tumor response to the drug of the respective resistant subline (Suppl. File 5). The expression of 91 genes was associated with patient response to two drugs, the expression of 51 genes associated with response to three drugs, the expression of 21 genes associated with expression to four drugs, and the expression of 6 genes associated with response to all five drugs (Fig. 4D, Suppl. File 5).

Considering the TNBC data alone, the expression of 165 genes was significantly correlated with patient survival in response to either doxorubicin, paclitaxel, or both drugs (Fig. 4E, Suppl. File 5). The expression of 141 of these 165 genes was associated with tumor response to the drug of the respective drug-adapted subline. The expression of 22 genes was associated with patient response to both doxorubicin and paclitaxel (Fig. 4E, Suppl. File 5).

Comparison of the analysis of the 165 genes identified in the TCGA analysis with the 81 genes identified in the analysis of *de novo* variants (Suppl. File 4) revealed 35 overlapping genes present in both datasets. This included 23 genes that have already been associated with drug resistance and 12 genes (*ABCD1, AGAP6, CUBN, DNAJC13, FLG, GXYLT1, KIAA0586, PABPC3, RGPD3, RGPD4, SETX and USP6*) that are novel findings (Suppl. File 6).

### Complex sensitivity patterns of drug-resistant sublines to cytotoxic drugs

Determining drug sensitivity profiles in the cell line panel against the drugs of adaptation, i.e., cisplatin, doxorubicin, eribulin, paclitaxel, gemcitabine, and 5-fluorouracil (Fig. 5A, Suppl. File 1), revealed complex resistance patterns that did not follow clear, predictable rules. For example, two of the three doxorubicin-adapted sublines (HCC38^r^DOX^40^ and HCC1806^r^DOX^12.5^) displayed increased (collateral) sensitivity to cisplatin compared to the parental cell line, while MDA-MB-468^r^DOX^50^ displayed cross-resistance to cisplatin (Fig. 5A, Suppl. File 1). Moreover, all resistant sublines remained sensitive to or showed collateral sensitivity to at least one of the other chemotherapeutic agents (Fig. 5A, Suppl. File 1). The 5-fluorouracil-resistant HCC1806^r^5-F^1500^ subline was the only resistant subline that remained sensitive to all other investigated cytotoxic drugs (Fig. 5A, Suppl. File 1).

**Figure 5.**
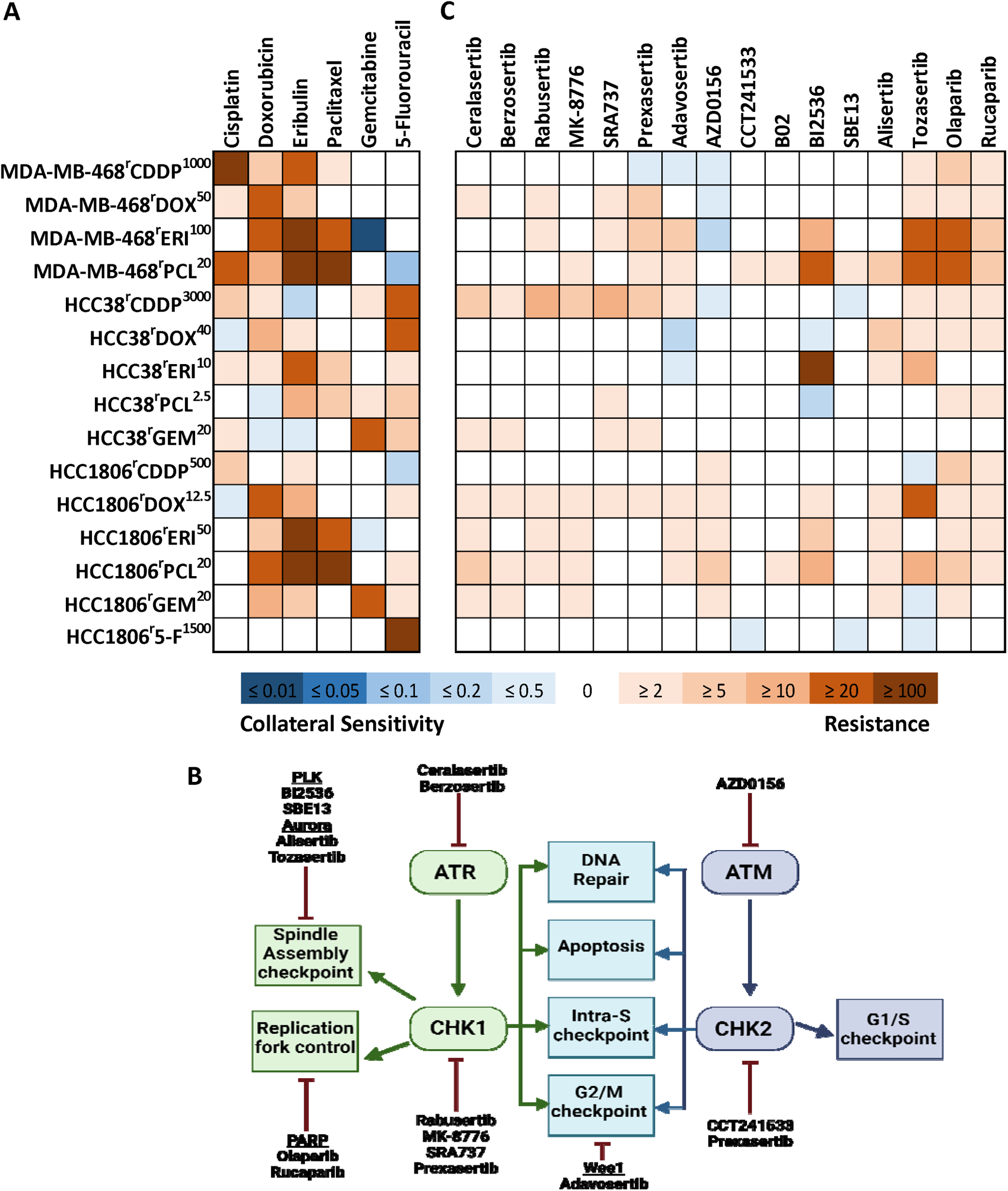
Complex sensitivity patterns to cytotoxic and DDR-targeted agents. A) Heatmap of fold resistance and collateral sensitivity to cytotoxic agents. B) Summary of pathways targeted by DNA damage response and repair (DDRR) inhibitors used in screening. C) Heatmap of fold change resistance and collateral sensitivity to DDRR inhibitors.

The ATP-binding cassette (ABC) transporter ABCB1 (also known as P-glycoprotein and MDR1) is an efflux transporter that mediates resistance to many anti-cancer drugs, including doxorubicin, eribulin, and paclitaxel ^46^. Only five of the nine sublines adapted to the ABCB1 substrates doxorubicin, eribulin, and paclitaxel (including all three eribulin-resistant sublines) displayed cross-resistance to all other ABCB1 substrates. Among the ABCB1 substrate-adapted sublines, all the eribulin-adapted sublines displayed cross-resistance to paclitaxel, and all the paclitaxel-adapted sublines displayed cross-resistance to eribulin (Fig. 5A, Suppl. File 1). Notably, eribulin and paclitaxel are both tubulin-binding agents but differ in their mechanisms of interaction with tubulin. Eribulin is a destabilizing agent that binds to the vinca binding site of tubulin and inhibits microtubule formation, while paclitaxel is a stabilizing agent that binds to the taxane binding site that impairs microtubule degradation ^47–51^. Further research will be required to determine to what extent the tubulin-binding agent cross-resistance profile of the tubulin-binding agent-adapted sublines is the consequence of the expression of ABCB1 (and/or other transporters), tubulin-related resistance mechanisms, or both.

Taken together, it is not possible to predict how resistance to a certain drug will affect the sensitivity patterns of the resulting sublines to other cytotoxic agents. However, all of the drug-resistant TNBC sublines remained sensitive and/or displayed collateral sensitivity to at least one of the tested anti-cancer drugs. Future research will be needed to elucidate the underlying mechanisms to identify biomarkers for personalized therapy approaches that can guide effective drugs to the right patients ^4^.

### Complex sensitivity patterns of drug-resistant sublines to DNA damage response (DDR) inhibitors

Triple-negative breast cancer cells have been shown to harbor defects in DNA damage repair signalling, which can result in a dependence on the remaining intact DNA damage repair (DDR) pathways and, in turn, in sensitivity to certain DDR inhibitors ^52^. Hence, we tested a panel of inhibitors targeting critical nodes of DDR signalling in our novel resistant TNBC cell line panel (Fig. 5B).

All parental cell lines displayed sensitivity to the tested DDR inhibitors at therapeutic concentrations, i.e., within the C_max_ values reported for these agents (for which this information was available) (Suppl. Fig. 7). Similar to the results obtained for the cytotoxic anti-cancer drugs, the DDR sensitivity profiles were complex and unpredictable in the resistant sublines (Fig. 5C, Suppl. File 1). Relative to the respective parental cell lines, the sensitivity remained unchanged for 128 DDR inhibitor/ resistant subline combinations. Increased resistance (cross-resistance) was detected in 96 DDR inhibitor/resistant subline combinations, and increased sensitivity (collateral vulnerability) was recorded in 16 DDR inhibitor/resistant subline combinations. Neither sublines of the same parental cell line nor sublines adapted to the same drugs displayed substantial overlap in their DDR inhibitor sensitivity profiles. Generally, cross-resistance levels were lowest for the CHK2 inhibitor CCT241533, the PLK1 inhibitor SBE13, and the RAD51 recombinase inhibitor B02 among the investigated DDR inhibitors (Fig. 5C, Suppl. File 1).

Cross-resistance patterns were even inconsistent between DDR inhibitors with the same targets. For example, different sensitivity patterns were observed between the ATR inhibitors ceralasertib and berzosertib as well as the CHK1 inhibitors rabusertib, MK-8776, SRA737, and prexasertib (Fig. 5C, Suppl. File 1). The reasons for these differences are unclear. Notably, the activity of the DDR inhibitors may be modified by interactions with additional targets, and off-target resistance mechanisms (e.g., processes associated with drug uptake or efflux) may contribute to these differences ^53^.

In summary, and in line with the findings from the investigation of cytotoxic anti-cancer drugs, the drug-adapted TNBC sublines displayed complex, unpredictable sensitivity patterns against DDR inhibitors. This further demonstrates that improved future therapies will depend on an advanced understanding of the underlying molecular processes that enable the identification of biomarkers that can guide effective therapies for individual patients after treatment failure^4^. Notably, CHK2, PLK1, and RAD51 may have potential as new drug targets for the discovery and development of next-line therapies for TNBC patients whose tumors have stopped responding to chemotherapy.

### Investigation of patterns in cell line drug response profiles

Finally, we used the delta (Δ) method to identify potential patterns in the response of the cell lines to all investigated cytotoxic anti-cancer drugs and DDR inhibitors ^54^. The IC_50_ values were transformed to ΔIC_50_ values for each compound (see methods) and correlated across the drug panel using linear regression analysis and testing for statistical significance (Suppl. Table 1). Positive correlations indicate that increased drug resistance is seen with both agents, whilst negative correlations indicate that whilst increasing drug resistance is observed for one agent, collateral sensitivity is observed for the other agent. In the MDA-MB-468, HCC38, and HCC1806 sublines, we observed 19, 20, and 60 positive correlations and 2, 8, and 1 negative correlation, respectively (Suppl. Table 1).

We were most interested in the agents that demonstrated negative correlations, as they may identify potential next-line treatments. However, among the 11 negative correlations, there were no consistent results across the cell line panel (Fig. 6). This further confirms that acquired resistance mechanisms are complex, individual, and unpredictable and that the identification of potential next-line therapies after treatment failure will depend on an improved understanding of cancer cell evolution enabling therapy monitoring and biomarker-guided treatment adaptation.

**Figure 6.**
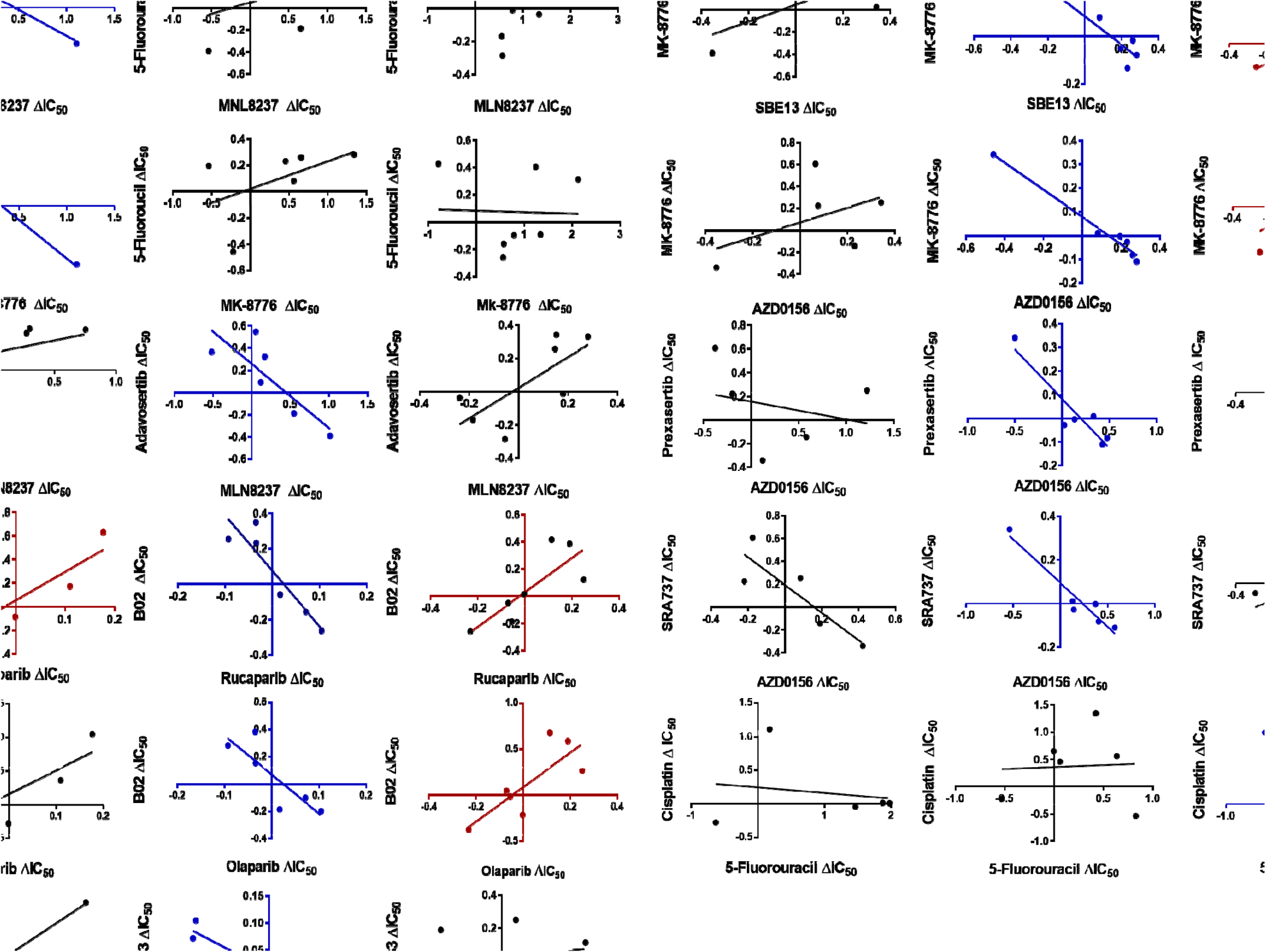
Lack of trends in drug or inhibitor sensitivity patterns. Graphs demonstrating a negative correlation; collateral sensitivity to one agent but resistance to the other (blue); positive correlation; resistance to both agents (red); and no statistical correlation (black) for each set of sublines adapted from the MDA-MB-468, HCC38 or HCC1806 TNBC cell lines.

## Discussion

In this study, we introduced and characterized a novel set of 15 sublines derived from the TNBC cell lines HCC38, HCC1806, and MDA-MB-468 that had been adapted to cisplatin, doxorubicin, eribulin, paclitaxel, gemcitabine, or 5-fluorouracil.

We applied whole exome sequencing to identify biomarker candidates to guide the use of anti-cancer therapies. In the first step, we focused on *de novo* mutations, i.e., mutations found in a resistant subline but undetectable in the respective parental cell line. Considering genes that displayed *de novo* mutations in at least two sublines of two different parental cell lines resulted in 81 resistance-associated variants, 48 of which were already known to be involved in cancer cell drug resistance, while 33 variants were novel.

In a second approach, we used TCGA data to investigate the potential clinical relevance of genes that harbored resistance-associated variants in the resistant sublines. In the pan-cancer dataset, the expression of 682 of these genes was correlated with patient survival in response to at least one of the investigated drugs. Considering only TNBC, the expression of 165 genes was significantly correlated with patient survival.

Comparison of the *de novo* variant analysis with the TNBC TCGA analysis identified 35 overlapping genes. Twenty-three of these genes are known to be associated with drug resistance. Twelve genes (*ABCD1, AGAP6, CUBN, DNAJC13, FLG, GXYLT1, KIAA0586, PABPC3, RGPD3, RGPD4, SETX and USP6*) are novel findings that may represent novel resistance biomarkers that have not been previously associated with drug resistance in cancer. Further research will be needed to investigate and define in more detail the role of these gene variants in cancer therapy response and the expression of these genes as biomarkers for the tailoring of personalized cancer therapies. Notably, numerous studies have shown that drug-adapted cancer cell lines exhibit clinically relevant resistance mechanisms^4,9–17^.

Interestingly, the analysis of exome sequencing data revealed remarkably few overlapping mutations between the investigated resistant sublines, including sublines derived from the same parental cell line and sublines adapted to the same drug. This suggests that resistance formation is the consequence of a complex, individual, and unpredictable evolutionary process.

This complexity was confirmed by the determination of drug sensitivity profiles to both cytotoxic anti-cancer drugs and DNA damage repair (DDR) inhibitors. Drug-adapted sublines of the same parental cell line and sublines adapted to the same drug displayed substantially different drug response patterns.

Notably, all the drug-adapted sublines remained sensitive and/or displayed increased sensitivity (collateral vulnerability) to a range of tested compounds. This suggests that it will be possible in the future to establish an improved understanding of the processes underlying acquired resistance formation that result in the identification of biomarkers that indicate effective next-line treatments for patients for whom currently no effective treatment is available.

Among the investigated DDR inhibitors, the CHK2 inhibitor CCT241533, the PLK1 inhibitor SBE13, and the RAD51 recombinase inhibitor B02 had the lowest cross-resistance levels. Thus, CHK2, PLK1, and RAD51 are potential drug targets in TNBC patients after failure of established therapies, particularly if reliable biomarkers are found that identify cancer patients who are likely to benefit from such treatments.

Overall, the results from the characterization of the project cell line panel indicated that cancer cell resistance is a complex, individual, and unpredictable process. This finding is in agreement with data from studies in which cancer cell lines were repeatedly adapted to the same drug in independent experiments ^8,16,55–57^ and with recent findings from a comprehensive analysis of cancer cell evolution in lung cancer patients ^58–62^.

In conclusion, we present a novel set of drug-adapted TNBC cell lines as preclinical models of acquired drug resistance. Overlapping genes detected through the characterization of *de novo* variants and patient-derived TCGA data identified 35 biomarker candidates for the guidance of personalized TNBC therapies for further investigation, including 12 novel genes that have not been previously associated with drug resistance in cancer. Finally, our results show that each cancer cell line adaptation process follows an individual, unpredictable route, which reflects recent clinical findings from the monitoring of cancer cell evolution in patients ^58–62^. This further supports the relevance of drug-adapted cancer cell lines as preclinical models of acquired resistance that can be analysed and manipulated at a level of detail that is impossible in the clinical setting.

## Materials and Methods

### Cell culture

MDA-MB-468, HCC38, and HCC1806 cells were obtained from the American Type Culture Collection (ATCC). The drug-adapted sublines (Fig. 1A, Suppl. File.1) were established by continuous exposure to stepwise increasing drug concentrations as previously described and derived from the Resistant Cancer Cell Line (RCCL) collection (https://research.kent.ac.uk/industrial-biotechnology-centre/the-resistant-cancer-cell-line-rccl-collection)^4,63^. All cell lines were cultured in Iscove’s Modified Dulbecco’s medium (IMDM) supplemented with 10% fetal bovine serum (Sigma□Aldrich, Germany), 2 mM L-glutamine, 25 mM HEPES (Fisher Scientific, UK), 100 IU/mL penicillin, and 100 µg/mL streptomycin (Life Technologies, UK) at 37 °C in a humidified atmosphere with 5% CO_2_. Each drug-adapted subline was continuously cultured in the presence of the specific adaptation drug at a defined concentration, as indicated by the cell line name (ng/mL), e.g., MDA-MB-468^r^DOX^50^, where r = the resistant subline, Dox = doxorubicin and 50 = 50 ng/ml.

### Compounds

The following compounds were purchased from the indicated suppliers: Adavosertib, Alisertib, Berzosertib, Ceralasertib, MK-8776, Olaparib, Prexasertib, Rabusertib, Rucaparib, SBE13, Tozasertib (Adooq Bioscience), AZD0156, BI2536, Doxorubicin, Gemcitabine (Selleckchem), B02, Cisplatin, 5-Fluorouracil (Sigma□Aldrich), CCT241533, SRA737 (a gift from the Institute of Cancer Research), Eribulin (Eisia), and Paclitaxel (Cayman Chemicals). All drug stocks were prepared in DMSO and stored at −20 °C, except for cisplatin, which was prepared in 0.9% NaCl solution and stored in the dark at room temperature.

### Cell growth and viability assays

Cell viability was tested using the 3-(4,5-dimethylthiazol-2-yl)-2,5-diphenyltetrazolium bromide (MTT) dye reduction assay after 120 hours of incubation with each compound, modified as previously described ^64,65^. Concentrations that reduced cell viability by 50% relative to an untreated control (IC_50_) were determined and used to calculate the resistance factor (RF; IC_50_ of drug-adapted cell line/IC_50_ of respective parental cell line).

### Whole exome sequencing

Whole exome sequencing (WES) libraries were prepared using the Nextera Rapid Capture Exome Kit (Illumina). Sequencing was performed on a HiSeq 1500 platform in Rapid Run mode with 2 x 100 nucleotide paired-end reads. The two lanes of the Rapid Run flow cell provided two sets of FASTQ data per cell line.

### Variant calling

FASTQC was used to control the quality of the raw sequence data ^66^ prior to the removal of sequencing adaptors. Trimmomatic (settings: NexteraPE-PE.fa:2:30:10 LEADING:3 TRAILING:3 SLIDING WINDOW: 4:15 MILEN:36) ^67^. Raw FASTQ files were aligned to the human reference genome (GRCH37) using Burrows□Wheeler Alignment (v.0.7.17) with an output in sequence alignment map (SAM) format applying the default settings -M -R ^68–70^. Only paired reads were used, and Samtools flagstat was used to print statistics throughout each of the subsequent steps ^68^. SAM files were input into Picard tools SortSam (v.2.17.10), where the read alignments were sorted by coordinate and converted to a binary alignment map (BAM) format (Picard Toolkit.2019. Broad Institute, GitHub Repository. http://broadinstitute.github.io/picard/; Broad Institute). Picard Tools MarkDuplicates (v2.17.10) was used for the removal of PCR duplicates (Picard Toolkit. 2019. Broad Institute, GitHub Repository. http://broadinstitute.github.io/picard/; Broad Institute). GenomeAnalysisTK-3.7.0 RealignerTargetCreator was used to perform base score recalibration, and GenomeAnalysisTK-3.7.0 IndelRealigner was used for INDEL realignment MAX_READS = 20000 ^71^. SAMtools mpileup was used to generate binary variant call format (BCF) files from the BAM files, which were then input into BCFtools to call the SNVs and INDELS to generate a variant calling format (VCF) ^72^. Variants were annotated with VEP ^73^.

### Variant filtering

Only variants in coding regions of the genome were considered. To identify high-confidence variants, variants with a Phred score < 30, variants with fewer than 10 reads supporting the base call, or variants with < 3 reads supporting the variant were removed. Moreover, common variants with a frequency of ≥ 0.001% in the genome aggregation database (gnomAD) were removed ^74^; if not, ≥ 3 samples were annotated in The Cancer Genome Atlas (TCGA), or ≥ 10 samples were annotated in the Catalogue Of Somatic Mutations In Cancer (COSMIC) ^39,75,76^.

### Definition of variants

*Gained* variants: variants that are called in the drug-resistant subline and are called with low confidence in the parental cell line. *De novo* variants: variants that are called in the drug-resistant subline but not called in the parental cell line. *Not called* variants: variants that are called in the parental cell line but not called in the drug-resistant subline, even at low confidence. *Lost* variants: variants that are called in the parental cell line and are called in low confidence in the drug-resistant subline. *Shared* variants: variants that are called in both the parental and drug-resistant sublines.

### Gene Ontology

Gene Ontology (GO) functional enrichment analysis was conducted using G:profiler ^77^. Gene lists were submitted as queries to the g:GOSt functional profiling tool and run at a significance threshold of g:SCS and a user threshold of 0.05.

### The Cancer Genome Atlas (TCGA) analysis

#### TCGA data retrieval

The data were collected from the UCSC Xena functional genomics browser [https://xenabrowser.net]. Batch-corrected gene expression data (RNAseq, log2(normalized value + 1)) for 11,060 patients (version 2016-12-29), clinical data for 12,591 patients (version 2018-09-13), and somatic mutation data (HG19) for 9,104 patients (version 2016-12-29) were downloaded for the TCGA pancancer (PANCAN) cohort. Curated drug data were obtained from Moiso 2021 for 4,321 patients ^78^.

#### Final datasets

The 4 downloaded datasets were filtered to a final dataset for each drug for which every data type was available (gene expression, somatic mutation, clinical, and drug data). If a patient did not have at least 1 somatic mutation recorded, they were excluded from further somatic mutation analyses. This resulted in final datasets of 683 patients (23 cancer types) treated with cisplatin, 385 (17) with doxorubicin, 367 (11) with fluorouracil, 349 (20) with gemcitabine, and 544 (16) with paclitaxel for which somatic mutation and clinical data were available (table – “mutations/treatment_by_cancer_type_mutation_patients.tsv”). The gene expression and treatment data included 765 patients (24 cancer types) treated with cisplatin, 571 (18) with doxorubicin, 452 (11) with fluorouracil, 438 (21) with gemcitabine, and 828 (15) with paclitaxel (table – “expression/treatment_by_cancer_type_expression_patients.tsv). Datasets including only TNBC patients were also created for further analysis^79^. This was only completed for those patients treated with doxorubicin (96 patients) and paclitaxel (63), as the number of patients treated with cisplatin (2), gemcitabine (4), and fluorouracil (21) was too low for meaningful analysis. For doxorubicin and paclitaxel treatments, gene expression and clinical data were available for 93 and 62 TNBC patients, respectively, while somatic mutation and clinical data were available for 74 doxorubicin- and 49 paclitaxel-treated patients. One TNBC patient (TCGA-AR-A256) whose disease-specific survival (DSS) data were incomplete was excluded.

#### Survival analysis

Analysis was performed in R version 4.3.0. Kaplan-Meier (KM) plots were generated for mutation status (mutated – MUT or wild type – WT) and for gene expression status (high or low) using the survival (v3.5-5) and survminer (0.4.9) packages. Somatic nonsynonymous mutations were considered in the genes of interest. The cut-off for high/low gene expression was calculated using the surv_cutpoint function in survminer, which makes use of the R package maxstat (v0.7-25). This gave a threshold for high/low expression based on the most significant relation with outcome, in this case, disease-specific survival. Any sample with gene expression > the calculated threshold was considered to have “high expression”, and any sample with gene expression < the threshold was considered to have “low expression”. The p value displayed on the KM plots was calculated using the log-rank test.

### Statistical analysis and data manipulation

GraphPad Prism 6 (GraphPad Software, Inc., USA) was used to generate dose□response curves and determine IC_50_ values via nonlinear regression (with variable slopes). Statistical significance was calculated using a two-tailed t-test, assuming unequal variance, in GraphPad Prism 6 (GraphPad Software, Inc., USA).

The delta method was used as described by Bracht *et al.*, 2006 ^54^. IC_50_ values were transformed to Δ IC_50_ values: Δ IC_50_ *= log (average* IC_50_ *in drug over all cell lines) – log (individual* IC_50_ *in drug for each cell line)*. Linear regression analysis of ΔIC_50X_ versus ΔIC_50Y,_ where X and Y represent two different compounds from the panel, was performed. The Pearson correlation coefficient (*r*) was used to establish the level of significance in a two-tailed test with (n-2) degrees of freedom, where p ≤ 0.05.

## Availability of data and materials

Data generated or analyzed during this study are included in this published article and its supplementary information files. The exome sequencing datasets generated and analyzed during the current study are available at Gene Expression Omnibus (GEO) repository (#PRJNA1155201).

## Competing interests

Nothing to declare.

## Funding

This work was supported by grants from the Frankfurter Stiftung für krebskranke Kinder, Kent Health, Kent Cancer Trust, and the Rosetrees Trust.

## Authors contribution

HEG, MMi, MDG, and MNW conceived and designed the study. HG, MA, IGR, KMM, AN, MMe, and JC acquired data. All authors analyzed and interpreted data. HEG, MM, MDG, and MNW drafted the work. All authors substantively revised the work and approved the submitted version.

## Acknowledgements

We would like to thank the Institute of Cancer Research for their kind gift of CCT241533 and SRA737. Figures were created using BioRender.com.

---

**Supplementary Figure 1. Chemo-naïve cell lines are clinically sensitive to chemotherapy agents.** IC_50_ values of drug-naïve parental cell lines treated with the respective chemotherapy agents: cisplatin, doxorubicin, eribulin, paclitaxel, gemcitabine or 5-fluorouracil. Green, MDA-MB-468 cells; blue, HCC38 cells; orange, HCC1806 cells. The black line indicates known C_max_ values for each chemotherapy agent. Data from n ≥ 3, statistics were calculated using Student’s t-test and are plotted as the mean ± SD.

**Supplementary Figure 2: Variant counts.** A) Total number of variants called for in the panel of drug-naïve and drug-resistant cell lines. B) Types of variants called for in the panel of drug-naïve and drug-resistant cell lines, including missense, synonymous, frameshift, inframe insertion, inframe deletion, stop loss, stop gain, splice acceptor and splice donor variants.

**Supplementary Figure 3. *De novo* variant overlaps.** The number of *de novo* variants that overlap in A) drug-resistant cell lines adapted to the same chemotherapy drug and B) drug-resistant cell lines adapted from the same parental cell line but to different chemotherapy drugs.

**Supplementary Figure 4. Gene ontology terms related to variants in drug-resistant sublines.** A) The number of variants increased in drug-resistant sublines (*de novo* variants, *gained* variants and *shared* variants that demonstrated a ≥2 increase in variant allele frequency). B) The number of variants decreased in drug-resistant sublines (*not-called* variants, *lost* variants and *shared* variants that demonstrated ≤2 decreases in variant allele frequency). The number and overlapping terms found in increased and decreased variants were compared between cell lines adapted to the same chemotherapy drug (C, E) and sublines derived from the same parental cell line but adapted to different chemotherapy drugs (D, F). Green bars indicate increased variants (A, C, D), and red bars indicate decreased variants (B, C, D).

**Supplementary Figure 5.** Chemo-naïve cell lines are clinically sensitive to DNA damage response and repair (DDRR) inhibitors. IC_50_ values of drug-naïve cell lines treated with the indicated drug. Green, MDA-MB-468-derived; blue, HCC38-derived; orange, HCC1806-derived. The black line indicates known C_max_ values for each DDRR agent. The data are from ≥ 3 independent experiments, and the statistics were calculated using Student’s t-test and are plotted as the means ± SDs.

**Supplementary Table 1. Drug correlation of delta (**Δ**) values.** The IC_50_ values were transformed to ΔIC_50_ values for each drug (see methods) and correlated across the drug panel, with linear regression analysis and statistical significance. The values in the table indicate the r values of the correlations, where positive values indicate positive correlations and negative values indicate negative correlations. P values of the correlations are indicated in the blue color scheme, with light blue (p≤0.05) indicating the lowest statistical significance and dark blue (p≤0.00001) indicating the highest statistical significance.

**Supplementary File 1.** Mean IC_50_ values, SDs and resistance factors for the project panel treated with chemotherapy drugs and DNA damage response inhibitors.

**Supplementary File 2.** Basic variant characterization of the cell line panel.

**Supplementary File 3.** Variants found to be *de novo*, *gained*, *not called*, *lost* and shared in drug-resistant cell lines.

**Supplementary File 4.** List of genes with *de novo* variants in ≥2 drug-resistant cell lines. The values in the table indicate the variant allele frequencies of the *de novo* variants identified in the indicated genes. PMIDs for genes previously implicated in cancer and drug resistance.

**Supplementary File 5.** Step-by-step analysis of both TNBC and pan-cancer patient data extracted from the TCGA.

**Supplementary File 6.** Comparison of genes identified through *de novo* variant analysis and TCGA analysis.

## Notes

### Competing Interest Statement

The authors have declared no competing interest.

### Summary of Updates

The TCGA analysis has been updated to a newer version.

