## Supplementary material for "Resistance patterns in drug-adapted cancer cell lines reflect the complex evolution in clinical tumours": Suppl Figures

### Supplementary Figure 1

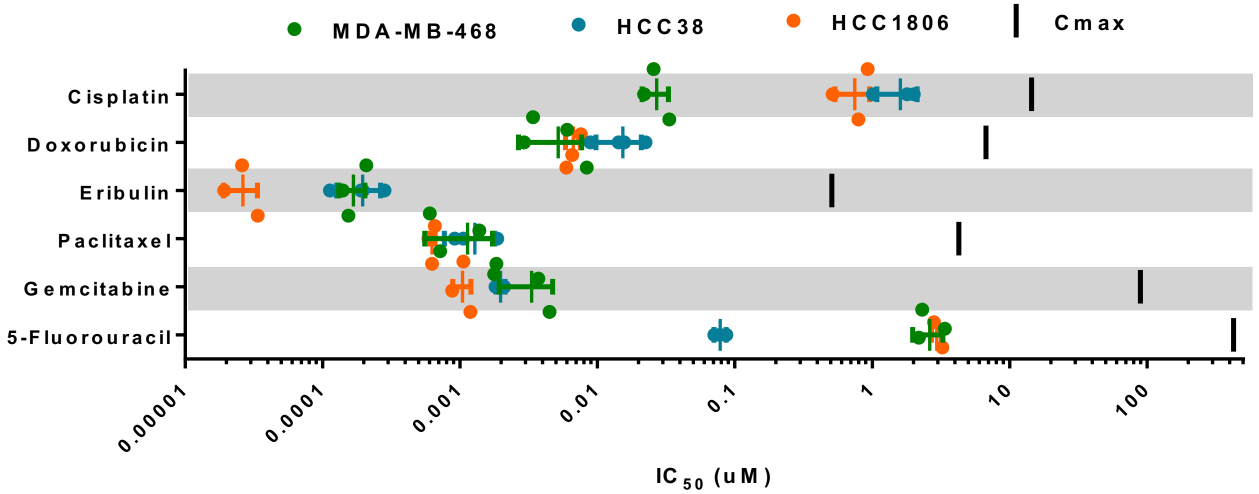


**Supplementary Figure 1. Chemo-naïve cell lines are clinically sensitive to chemotherapy agents.** $IC_{50}$ values of drug-naïve parental cell lines treated with the respective chemotherapy agents: cisplatin, doxorubicin, eribulin, paclitaxel, gemcitabine or 5-fluorouracil. Green, MDA-MB-468 cells; blue, HCC38 cells; orange, HCC1806 cells. The black line indicates known $C_{max}$ values for each cytotoxic drug. Data from $n \geq 3$, statistics were calculated using Student's t-test and are plotted as the mean ± SD.

### Supplementary Figure 2

A

B

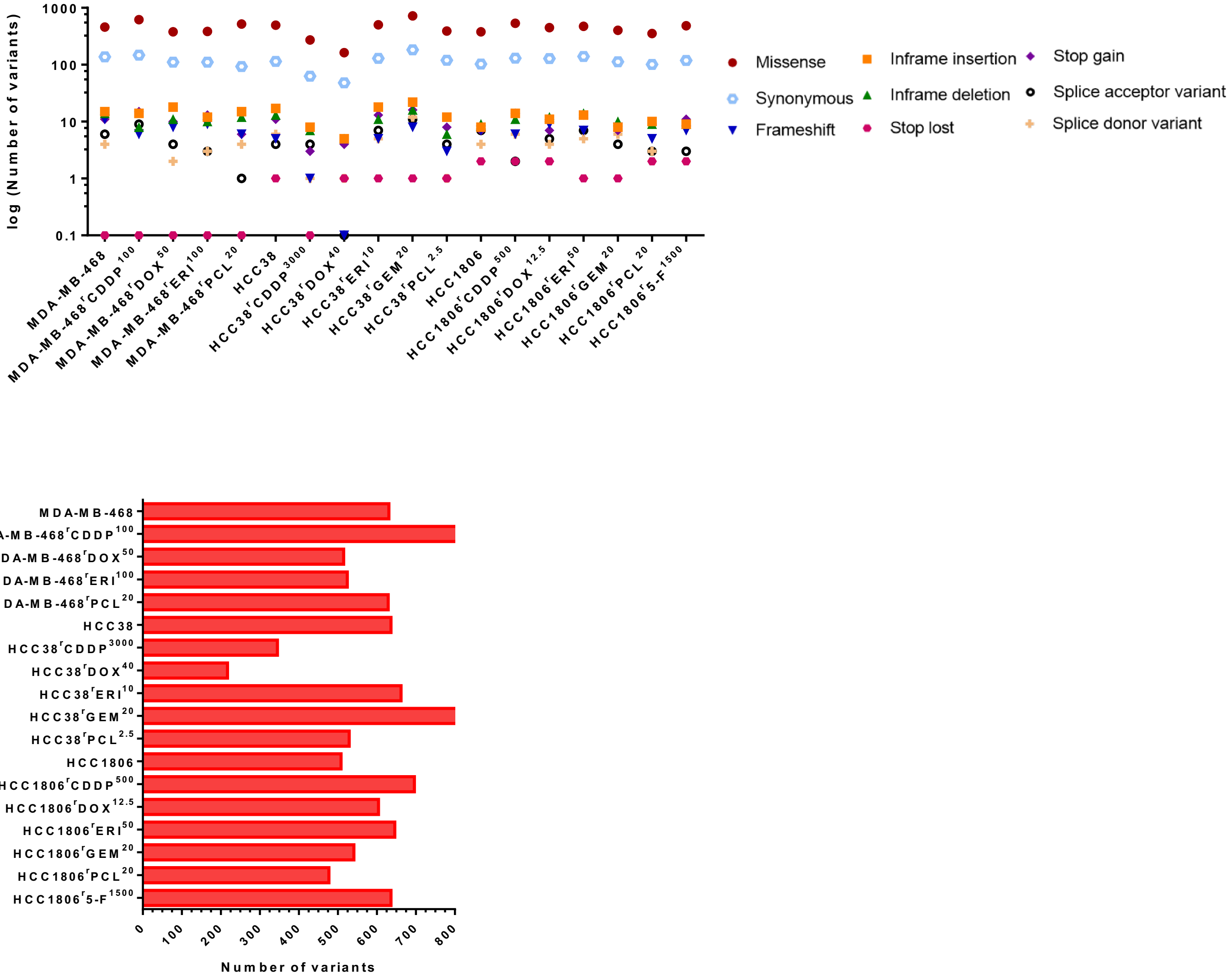


**Supplementary Figure 2. Variant counts.** A) Total number of variants called for in the panel of drug-naïve and drug-resistant cell lines. B) Types of variants called for in the panel of drug-naïve and drug-resistant cell lines, including missense, synonymous, frameshift, inframe insertion, inframe deletion, stop loss, stop gain, splice acceptor and splice donor variants.

### Supplementary Figure 3

**A**

**Cisplatin**

| | MDA-MB-468 | HCC38 | HCC1806 |
|---|---|---|---|
| **MDA-MB-468** | 213 | | |
| **HCC38** | 0 | 98 | |
| **HCC1806** | 1 | 0 | 155 |

**Doxorubicin**

| | MDA-MB-468 | HCC38 | HCC1806 |
|---|---|---|---|
| **MDA-MB-468** | 79 | | |
| **HCC38** | 0 | 31 | |
| **HCC1806** | 0 | 4 | 98 |

**Eribulin**

| | MDA-MB-468 | HCC38 | HCC1806 |
|---|---|---|---|
| **MDA-MB-468** | 89 | | |
| **HCC38** | 6 | 131 | |
| **HCC1806** | 5 | 3 | 121 |

**Paclitaxel**

| | MDA-MB-468 | HCC38 | HCC1806 |
|---|---|---|---|
| **MDA-MB-468** | 225 | | |
| **HCC38** | 2 | 92 | |
| **HCC1806** | 3 | 0 | 131 |

**Gemcitabine**

| | HCC38 | HCC1806 |
|---|---|---|
| **HCC38** | 203 | |
| **HCC1806** | 1 | 88 |

**B**

| | MDA-MB-468$^r$CDDP$^{1000}$ | MDA-MB-468$^r$DOX$^{50}$ | MDA-MB-468$^r$ERI$^{50}$ | MDA-MB-468$^r$PCL$^{20}$ |
|---|---|---|---|---|
| **MDA-MB-468$^r$CDDP$^{1000}$** | 213 | | | |
| **MDA-MB-468$^r$DOX$^{50}$** | 18 | 79 | | |
| **MDA-MB-468$^r$ERI$^{50}$** | 16 | 6 | 89 | |
| **MDA-MB-468$^r$PCL$^{20}$** | 5 | 6 | 4 | 225 |

| | HCC38$^r$CDDP$^{3000}$ | HCC38$^r$DOX$^{40}$ | HCC38$^r$ERI$^{10}$ | HCC38$^r$PCL$^{2.5}$ | HCC38$^r$GEM$^{20}$ |
|---|---|---|---|---|---|
| **HCC38$^r$CDDP$^{3000}$** | 98 | | | | |
| **HCC38$^r$DOX$^{40}$** | 3 | 31 | | | |
| **HHCC38$^r$ERI$^{10}$** | 11 | 9 | 131 | | |
| **HCC38$^r$PCL$^{2.5}$** | 3 | 2 | 19 | 92 | |
| **HCC38$^r$GEM$^{20}$** | 8 | 16 | 53 | 27 | 203 |

| | HCC1806$^r$CDDP$^{500}$ | HCC1806$^r$DOX$^{12.5}$ | HCC1806$^r$ERI$^{50}$ | HCC1806$^r$PCL$^{20}$ | HCC1806$^r$GEM$^{20}$ | HCC1806$^r$5-F$^{1500}$ |
|---|---|---|---|---|---|---|
| **HCC1806$^r$CDDP$^{500}$** | 155 | | | | | |
| **HCC1806$^r$DOX$^{12.5}$** | 27 | 98 | | | | |
| **HCC1806$^r$ERI$^{50}$** | 30 | 28 | 121 | | | |
| **HCC1806$^r$PCL$^{20}$** | 24 | 17 | 25 | 64 | | |
| **HCC1806$^r$GEM$^{20}$** | 19 | 22 | 31 | 17 | 88 | |
| **HCC1806$^r$5-F$^{1500}$** | 20 | 23 | 30 | 19 | 21 | 131 |

**Supplementary Figure 3: *De novo* variant overlaps.** The number of *de novo* variants that overlap in A) drug-resistant cell lines adapted to the same chemotherapy drug and B) drug-resistant cell lines adapted from the same parental cell line but to different chemotherapy drugs.

### Supplementary Figure 4

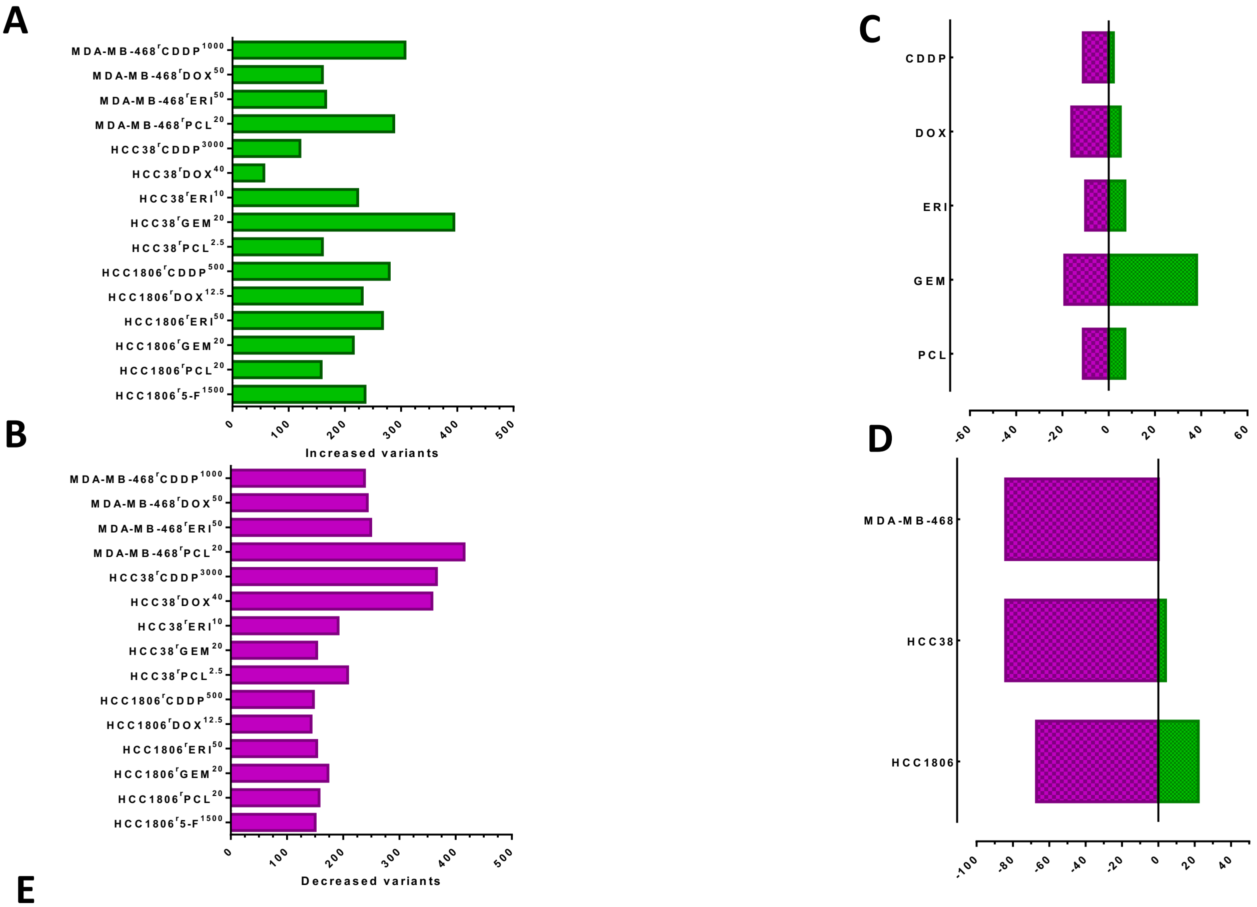


## E

| | Increased | | | Decreased | | |
|---|---|---|---|---|---|---|
| CDDP | GO:CC | GO:0070701 | mucus layer | GO:MF | GO:0030197 | extracellular matrix constituent, lubricant activity |
| | | | | GO:BP | GO:0030277 | maintenance of gastrointestinal epithelium |
| | | | | GO:CC | GO:0005796 | Golgi lumen |
| DOX | GO:BP | GO:0030277 | maintenance of gastrointestinal epithelium | GO:MF | GO:0008266 | poly(U) RNA binding |
| | GO:CC | GO:0005796 | Golgi lumen | GO:MF | GO:0070717 | poly-purine tract binding |
| | | | | GO:BP | GO:0015990 | electron transport coupled proton transport |
| | | | | GO:CC | GO:0005796 | Golgi lumen |
| ERI | GO:BP | GO:0030277 | maintenance of gastrointestinal epithelium | GO:MF | GO:0030197 | extracellular matrix constituent, lubricant activity |
| | | | | GO:BP | GO:0030277 | maintenance of gastrointestinal epithelium |
| | | | | GO:CC | GO:0005796 | Golgi lumen |
| GEM | GO:MF | GO:0030197 | extracellular matrix constituent, lubricant activity | GO:MF | GO:0030197 | extracellular matrix constituent, lubricant activity |
| | GO:CC | GO:0005796 | Golgi lumen | GO:CC | GO:0005796 | Golgi lumen |
| | | | | GO:CC | GO:0031012 | extracellular matrix |
| | | | | GO:CC | GO:0005680 | anaphase-promoting complex |
| PCL | GO:MF | GO:0005201 | extracellular matrix structural constituent | GO:MF | GO:0005201 | extracellular matrix structural constituent |
| | GO:BP | GO:0030277 | maintenance of gastrointestinal epithelium | GO:BP | GO:0030277 | maintenance of gastrointestinal epithelium |
| | GO:CC | GO:0005796 | Golgi lumen | GO:CC | GO:0005796 | Golgi lumen |
| | | | | GO:CC | GO:0031012 | extracellular matrix |

## F

| | Increased | | | Decreased | | |
|---|---|---|---|---|---|---|
| MDA-MB-468 | | | | GO:MF | GO:0030197 | extracellular matrix constituent, lubricant activity |
| | | | | GO:CC | GO:0016020 | membrane |
| | | | | GO:CC | GO:0005796 | Golgi lumen |
| | | | | GO:CC | GO:0097381 | photoreceptor disc membrane |
| HCC38 | | | | GO:MF | GO:0030197 | extracellular matrix constituent, lubricant activity |
| HCC1806 | GO:MF | GO:0030197 | extracellular matrix constituent, lubricant activity | GO:CC | GO:0005796 | Golgi lumen |
| | GO:MF | GO:0003727 | single-stranded RNA binding | | | |
| | GO:CC | GO:0005796 | Golgi lumen | | | |

### Supplementary Figure 4

**Supplementary Figure 4. Gene ontology terms related to variants in drug-resistant sublines.** A) The number of variants increased in drug-resistant sublines (*de novo* variants, *gained* variants and *shared* variants that demonstrated a ≥2 increase in variant allele frequency). B) The number of variants decreased in drug-resistant sublines (*not-called* variants, *lost* variants and *shared* variants that demonstrated ≤2 decreases in variant allele frequency). The number and overlapping terms found in increased and decreased variants were compared between cell lines adapted to the same chemotherapy drug (C, E) and sublines derived from the same parental cell line but adapted to different chemotherapy drugs (D, F). Green bars indicate increased variants (A, C, D), and red bars indicate decreased variants (B, C, D).

### Supplementary Figure 5

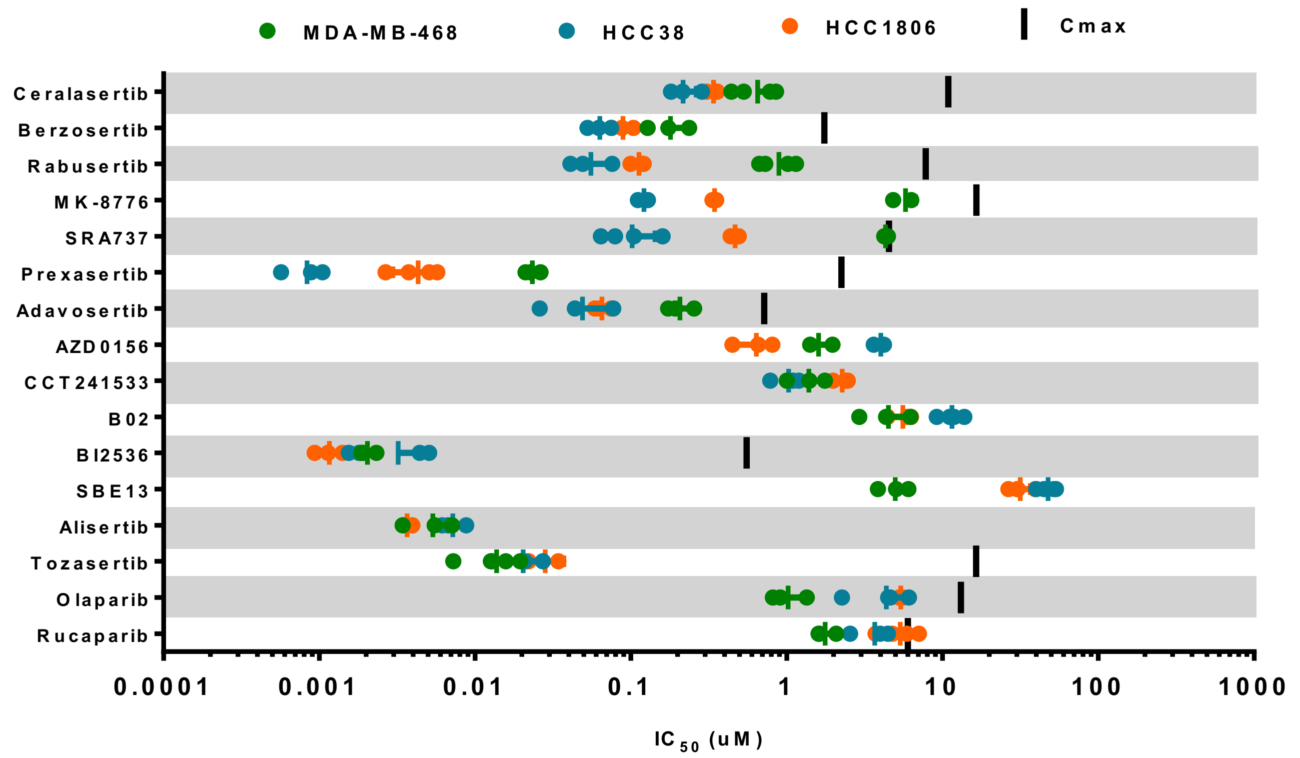


**Supplementary Figure 5. Chemo-naïve cell lines are clinically sensitive to DNA damage response and repair (DDRR) inhibitors.** $IC_{50}$ values of drug-naïve cell lines treated with the indicated drug. Green, MDA-MB-468-derived; blue, HCC38-derived; orange, HCC1806-derived. The black line indicates known $C_{max}$ values for each DDRR agent. The data are from ≥ 3 independent experiments, and the statistics were calculated using Student's t-test and are plotted as the means ± SDs.
