## Supplementary material for "Resistance patterns in drug-adapted cancer cell lines reflect the complex evolution in clinical tumours": Suppl Table 1

**Supplementary Table 1: Drug correlation of delta (Δ) values.** The $IC_{50}$ values were transformed to $\Delta IC_{50}$ values for each drug (see methods) and correlated across the drug panel, with linear regression analysis and statistical significance. The values in the table indicate the r values of the correlations, where positive values indicate positive correlations and negative values indicate negative correlations. P values of the correlations are indicated in the blue color scheme, with light blue ($p \leq 0.05$) indicating the lowest statistical significance and dark blue ($p \leq 0.00001$) indicating the highest statistical significance.

| | | | | | Microtubule | | Chk1 | | | Chk1/Chk2 | ATR | | Wee1 | ATM | CHK2 | RAD51 | PLK1 | PLK1,2 | Aurora A | Aurora (pan) | PARP | |
|---|---|---|---|---|---|---|---|---|---|---|---|---|---|---|---|---|---|---|---|---|---|---|
| | CDDP | DOX | GEM | 5-FU | ERI | PCL | Rabusertib | MK-8776 | SRA737 | Prexasertib | Ceralasertib | Berzosertib | Adavosertib | AZD0156 | CCT241533 | B02 | BI2536 | SBE13 | Alisertib | Tozasertib | Olaparib | Rucaparib |
| **CDDP** | 1 | -0.2163 | 0.4467 | -0.2803 | -0.1521 | -0.1925 | -0.3598 | -0.4702 | -0.4803 | -0.6388 | -0.04013 | -0.1964 | -0.4139 | 0.3788 | -0.1229 | 0.5292 | 0.1183 | 0.5312 | 0.0758 | -0.1047 | 0.2464 | 0.2409 |
| **DOX** | | 1 | -0.2533 | 0.218 | 0.4911 | 0.4247 | 0.3333 | 0.367 | 0.2748 | 0.4056 | 0.4732 | 0.3907 | 0.1615 | 0.03466 | 0.3736 | 0.102 | 0.3238 | 0.0006027 | 0.548 | 0.7374 | 0.5101 | 0.6071 |
| **GEM** | | | 1 | -0.1957 | -0.3474 | -0.4456 | -0.2164 | -0.2231 | -0.2579 | -0.3277 | 0.06139 | 0.004831 | -0.3523 | 0.4624 | -0.1891 | 0.003953 | -0.1257 | 0.265 | 0.03104 | -0.334 | -0.2629 | -0.1892 |
| **5-FU** | | | | 1 | -0.1574 | -0.1287 | 0.2138 | 0.1163 | 0.1678 | 0.1923 | 0.1211 | 0.07552 | 0.0637 | -0.4094 | -0.03066 | -0.3363 | -0.2507 | -0.2026 | -0.2481 | 0.1578 | -0.01432 | 0.1692 |
| **ERI** | | | | | 1 | 0.9316 | 0.2646 | 0.5057 | 0.2759 | 0.3515 | 0.4576 | 0.28 | 0.3736 | 0.09283 | 0.5255 | 0.3497 | 0.4849 | 0.003682 | 0.5302 | 0.5732 | 0.4877 | 0.432 |
| **PCL** | | | | | | 1 | 0.2828 | 0.5044 | 0.3011 | 0.4191 | 0.4632 | 0.3191 | 0.4968 | 0.0151 | 0.536 | 0.3396 | 0.461 | -0.04814 | 0.4678 | 0.5755 | 0.5048 | 0.4141 |
| **Rabusertib** | | | | | | | 1 | 0.8622 | 0.9423 | 0.8495 | 0.7718 | 0.8569 | 0.7768 | -0.587 | 0.5798 | -0.383 | -0.02318 | -0.7504 | -0.2507 | 0.1571 | -0.03476 | 0.02465 |
| **MK-8776** | | | | | | | | 1 | 0.9066 | 0.9075 | 0.725 | 0.7601 | 0.7748 | -0.4483 | 0.7199 | -0.4066 | 0.04952 | -0.6823 | 0.0809 | 0.2715 | 0.05423 | 0.1019 |
| **SRA737** | | | | | | | | | 1 | 0.8908 | 0.6405 | 0.7653 | 0.7375 | -0.6483 | 0.6167 | -0.4938 | -0.04883 | -0.8533 | -0.1694 | 0.1244 | -0.08551 | -0.04682 |
| **Prexasertib** | | | | | | | | | | 1 | 0.6641 | 0.7574 | 0.821 | -0.5751 | 0.6338 | -0.481 | 0.006958 | -0.7168 | -0.03754 | 0.25 | 0.04151 | 0.06158 |
| **Ceralasertib** | | | | | | | | | | | 1 | 0.9282 | 0.7152 | -0.1329 | 0.5623 | 0.02164 | 0.1873 | -0.266 | 0.03747 | 0.4642 | 0.3707 | 0.4091 |
| **Berzosertib** | | | | | | | | | | | | 1 | 0.7955 | -0.2974 | 0.4671 | -0.1797 | 0.1749 | -0.4767 | -0.1419 | 0.3467 | 0.1682 | 0.1727 |
| **Adavosertib** | | | | | | | | | | | | | 1 | -0.4665 | 0.4403 | -0.1911 | 0.02962 | -0.5047 | -0.217 | 0.2699 | 0.1571 | 0.1153 |
| **AZD0156** | | | | | | | | | | | | | | 1 | -0.3187 | 0.6437 | 0.1868 | 0.7096 | 0.5452 | 0.1791 | 0.1962 | 0.1908 |
| **CCT241533** | | | | | | | | | | | | | | | 1 | -0.1923 | -0.02043 | -0.3443 | 0.2498 | 0.3135 | 0.2793 | 0.3418 |
| **B02** | | | | | | | | | | | | | | | | 1 | 0.318 | 0.6656 | 0.3407 | 0.4032 | 0.4972 | 0.4147 |
| **BI2536** | | | | | | | | | | | | | | | | | 1 | 0.1033 | 0.413 | 0.4083 | 0.104 | -0.01209 |
| **SBE13** | | | | | | | | | | | | | | | | | | 1 | 0.3437 | 0.2763 | 0.4127 | 0.4117 |
| **Alisertib** | | | | | | | | | | | | | | | | | | | 1 | 0.5358 | 0.4058 | 0.4593 |
| **Tozasertib** | | | | | | | | | | | | | | | | | | | | 1 | 0.6121 | 0.6714 |
| **Olaparib** | | | | | | | | | | | | | | | | | | | | | 1 | 0.9433 |
| **Rucaparib** | | | | | | | | | | | | | | | | | | | | | | 1 |
